# Chikungunya virus infection disrupts MHC-I antigen presentation via nonstructural protein 2

**DOI:** 10.1101/2023.11.03.565436

**Authors:** Brian C. Ware, M. Guston Parks, Thomas E. Morrison

**Affiliations:** Department of Immunology and Microbiology, University of Colorado Anschutz Medical Campus, Aurora, Colorado, USA

## Abstract

Infection by chikungunya virus (CHIKV), a mosquito-borne alphavirus, causes severe polyarthralgia and polymyalgia, which can last in some people for months to years. Chronic CHIKV disease signs and symptoms are associated with the persistence of viral nucleic acid and antigen in tissues. Like humans and nonhuman primates, CHIKV infection in mice results in the development of robust adaptive antiviral immune responses. Despite this, joint tissue fibroblasts survive CHIKV infection and can support persistent viral replication, suggesting that they escape immune surveillance. Here, using a recombinant CHIKV strain encoding a chimeric protein of VENUS fused to a CD8^+^ T cell epitope, SIINFEKL, we observed a marked loss of both MHC class I (MHC-I) surface expression and antigen presentation by CHIKV-infected joint tissue fibroblasts. Both *in vivo* and *ex vivo* infected joint tissue fibroblasts displayed reduced cell surface levels of H2-K^b^ and H2-D^b^ MHC proteins while maintaining similar levels of other cell surface proteins. Mutations within the methyl transferase-like domain of the CHIKV nonstructural protein 2 (nsP2) increased MHC-I cell surface expression and antigen presentation efficiency by CHIKV-infected cells. Moreover, expression of WT nsP2 alone, but not nsP2 with mutations in the methyltransferase-like domain, resulted in decreased MHC-I antigen presentation efficiency. MHC-I surface expression and antigen presentation could be rescued by replacing VENUS-SIINFEKL with SIINFEKL tethered to β2-microglobulin in the CHIKV genome, which bypasses the need for peptide processing and TAP-mediated peptide transport into the endoplasmic reticulum. Collectively, this work suggests that CHIKV escapes the surveillance of antiviral CD8^+^ T cells, in part, by nsP2-mediated disruption of MHC-I antigen presentation.

**AUTHOR SUMMARY:** Arthritogenic alphaviruses, including chikungunya virus (CHIKV), are re-emerging global public health threats with no approved vaccines or antiviral therapies. Infection with these viruses causes debilitating musculoskeletal disease for months to years that is associated with the persistence of viral RNA and antigen. Prior studies using a mouse model found that CD8^+^ T cells, which recognize viral peptides in the context of major histocompatibility class I (MHC-I) displayed on the surface of infected cells, have a limited role in the control and clearance of CHIKV infection in joint-associated tissues, suggesting that CHIKV infected cells evade these critical effectors of the anti-viral immune response. Here, we show that MHC-I antigen presentation is inefficient in CHIKV-infected joint tissue fibroblasts, and that a protein encoded by CHIKV and produced in infected cells, nonstructural protein 2 (nsP2), disrupts the surface display of MHC-I molecules and antigen recognition of infected cells by CD8^+^ T cells. Our findings support a role for CHIKV nsP2 in the evasion of the CD8^+^ T cell response and establishment of persistent infection.

## INTRODUCTION

Since its re-emergence in 2004 in the Indian Ocean region (1, 2), chikungunya virus (CHIKV) has spread to numerous subtropical and tropical regions of Asia (3), the Caribbean (4), the Americas (2) and Europe (5) causing large-scale mosquito-borne epidemics that challenge public health systems. In the Americas, over three million CHIKV cases in 45 countries have been documented, and the global spread of CHIKV places approximately 1.3 billion people at risk for infection (6). CHIKV infection causes an acute debilitating febrile illness accompanied by high fever, and severe myalgia, and arthralgia. Musculoskeletal tissue pain and inflammation can become chronic in up to two thirds of patients (7, 8, 9). Due to the increasing incidence of CHIKV disease, the unpredictable nature of its outbreaks, and the lack of approved vaccines or anti-viral medications, it remains critical to elucidate CHIKV pathogenesis and immune restriction and evasion mechanisms.

CHIKV is one of ∼32 *Togaviridae* family members (10) and is comprised of an 11.8 kilobase, positive-sense, single-stranded RNA genome that contains two open reading frames encoding nonstructural and structural polyproteins (11). The initial viral RNA (vRNA) transcript enters the cytoplasm after viral binding and fusion, and two viral nonstructural polyproteins termed P123 and P1234 are translated (11). The protease activity of nonstructural protein 2 (nsP2) cleaves P1234 into P123 and nsP4, thus allowing for formation of a replication complex that synthesizes full-length negative-strand RNA from the template viral genome. Subsequent processing of P123 into nsP1, nsP2, and nsP3 results in formation of a replicase that synthesizes positive sense genomic vRNA as well as a subgenomic mRNA that is translated into the viral structural polyprotein. Though nsP1-4 are synthesized at equal molar ratios in their polyprotein form, only a single nsP2 for every twelve nsP1s is found within the replicase complex (12, 13), suggesting nsP2 has auxiliary functions apart from the replicase. Indeed, nsP2 has been reported to enter nuclei (14), suppress host mRNA transcription through the induction of RPB1 ubiquitination and degradation (15), disrupt type I interferon (IFN) signaling (16, 17), and suppress the unfolded protein response (UPR) (18).

CD8^+^ T cells are key executors of the adaptive immune response against many viral infections, including some arbovirus infections (19, 20, 21, 22, 23, 24). Their capacity to surveil, recognize, and lyse virus-infected cells is well documented (25, 26, 27, 28). Early innate immune responses, including inflammatory cytokines (29, 30), and the presence of viral peptides displayed by major histocompatibility complex class I (MHC-I) molecules on cell surfaces dictate the breadth and duration of the CD8^+^ T cell response. Importantly, CD8^+^ T cells scan for the presence of viral peptide-loaded MHC-I (pMHC-I) complexes on the surface of infected cells via their T cell receptor (TCR) (31, 32). Ligation of the TCR leads to activation and release of cytotoxic effectors such as perforin 1 and granzyme B and the subsequent destruction of the infected target cell (33). Numerous viruses have evolved mechanisms to subvert this crucial immune surveillance system through downregulation, suppression, or elimination of pMHC-I complexes from the surface of infected cells, thereby evading CD8^+^ T cell-mediated clearance (34). Although this paradigm has been extensively illustrated for DNA viruses such as herpesviruses and poxviruses (35), some RNA viruses also have evolved mechanisms for interfering with CD8^+^ T cell recognition. For example, the human immunodeficiency virus 1 Nef protein diverts pMHC-I complexes to the protein degradation pathway (36, 37), and infection with influenza A and B viruses also suppresses surface MHC-I expression (38). SARS-CoV-2 infection was shown to disrupt MHC-I cell surface expression (39, 40), in part, by disrupting nuclear transport of NLRC5 (40), the principal MHC-I transcription factor.

Recently, we observed that during CHIKV infection, CD8^+^ T cells accumulate in joint-associated tissue and lyse exogenously peptide-loaded splenocytes that had been adoptively transferred into the tissue. Despite this finding, CHIKV-specific CD8^+^ T cells appear to have little to no impact on the clearance of CHIKV-infected cells, as the viral burden in joint tissues of WT and *Cd8α^-/-^* mice are similar throughout the course of infection, suggesting that CHIKV infected cells evade the CD8^+^ T cell response (41).

Here, using a recombinant chimeric protein composed of the reporter VENUS fused to a well characterized CD8^+^ T cell epitope (ovalbumin 257-264; SIINFEKL), imbedded in the CHIKV genome, we observed a marked loss of both MHC-I cell surface expression and antigen presentation by CHIKV-infected joint tissue fibroblasts. Mutations within the methyl transferase-like domain of the CHIKV nsP2 restored MHC-I cell surface expression and antigen presentation efficiency by CHIKV-infected cells. In contrast to cell surface MHC-I expression, intracellular MHC-I levels were unaffected by CHIKV infection. Moreover, MHC-I antigen presentation by CHIKV-infected cells was rescued by replacing the VENUS-SIINFEKL in the CHIKV genome with SIINFEKL tethered to β2-microglobulin(β_2_m), which obviates the need for peptide processing and TAP-mediated peptide transport into the endoplasmic reticulum. Finally, we find that expression of wild-type (WT) nsP2 alone, but not nsP2 with mutations in the methyltransferase-like domain, is sufficient to diminish MHC-I antigen presentation efficiency in primary joint tissue fibroblasts. Collectively, this work suggests that CHIKV-infected cells escape surveillance by antiviral CD8^+^ T cells, in part, by nsP2-mediated disruption of MHC-I antigen presentation.

## RESULTS

### CHIKV infected primary joint tissue fibroblasts display decreased MHC-I cell surface expression

To determine if virus-encoded peptides are presented by MHC-I on CHIKV-infected cells, a recombinant CHIKV strain (CHIKV-VENKL) was engineered to express the chimeric protein VENUS-SIINFEKL (VENKL) wherein VENUS, a fluorescent reporter protein (42), and the CD8^+^ T cell peptide SIINFEKL from ovalbumin (43) were encoded between the capsid autoproteolysis domain (44) and an engineered 2a ribosomal skip site in-frame in the structural open reading frame of the CHIKV genome (**Fig 1A**), a design we have used previously (45). SIINFEKL follows the N-terminus of VENUS flanked by upstream (LELQE) and downstream (TEW) amino acid residues important for efficient peptide processing and MHC-I loading of SIINFEKL (46). This design allows for the measurement of MHC-I antigen presentation efficiency because we can measure SIINFEKL presentation as a function of source protein (i.e., VENUS) across all cells on a per-cell basis (46). To assess MHC-I antigen presentation by physiologically relevant CHIKV target cells, we generated primary *ex vivo* ankle (EVA) cell cultures by adapting a previously established protocol in which single cell suspensions were generated from murine ankle tissue (41), depleted of erythrocytes and nonadherent cells, and then cultured in growth medium.

**Figure 1.**
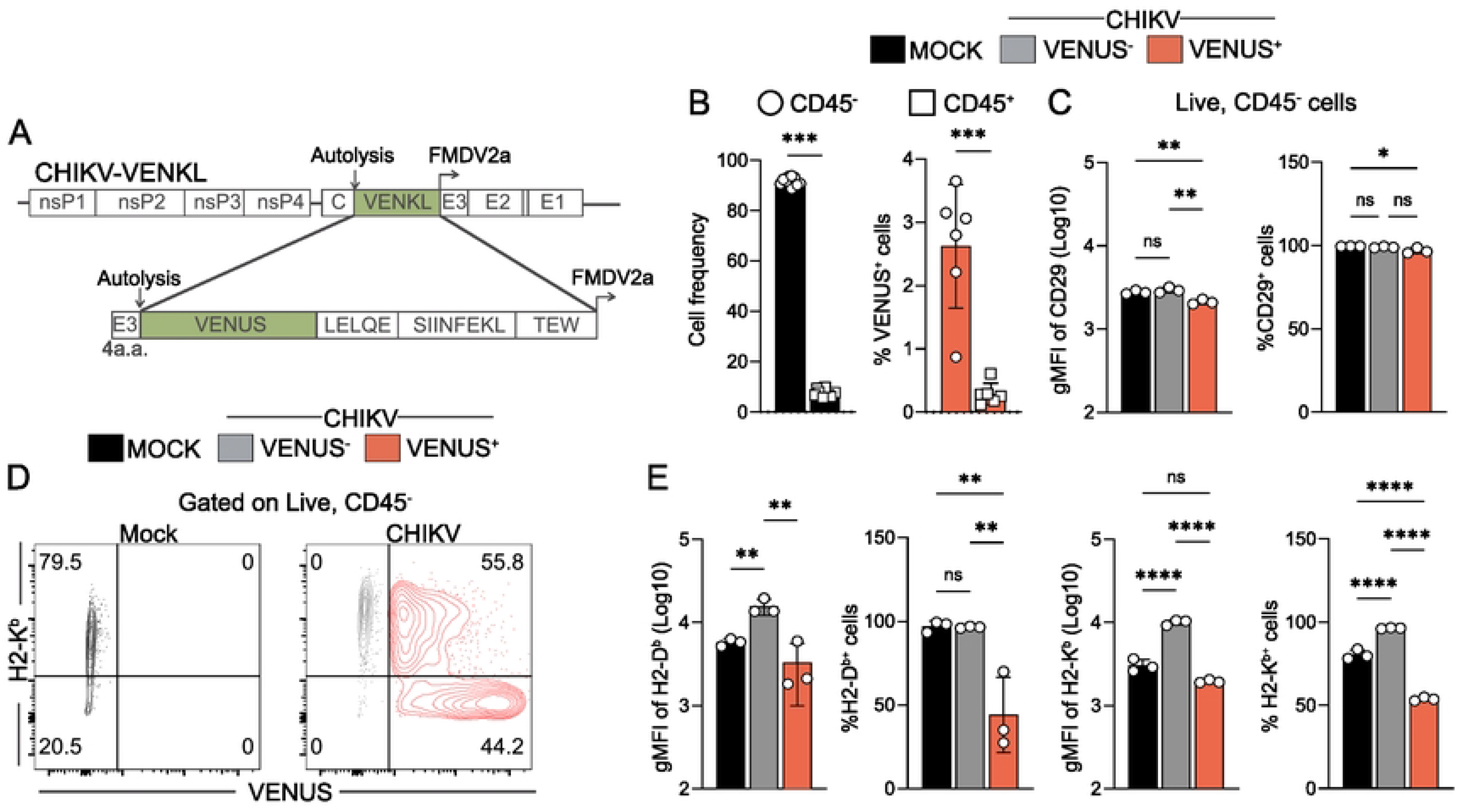
CHIKV infected primary joint tissue fibroblasts display decreased MHC-I cell surface expression. (**A**) Schematic of the recombinant CHIKV-VENKL. The coding sequence for the VENUS-SIINFEKL chimeric protein was inserted in-frame into the structural ORF of the CHIKV genome with known cleavage motifs flanking SIINFEKL. (**B-E**) EVA cells were inoculated with PBS (mock) or 10^6^ PFU CHIKV-VENKL. (**B**) At 24 hpi, the percentages of CD45^-^ (open circles) and CD45^+^ (open squares) cells among mock-inoculated cells and VENUS^+^ cells were determined. (**C**) The magnitude and frequency of expression of CD29 on CD45^-^ cells was determined by flow cytometry. (**D**) Representative flow cytometry plots of H2-K^b^ and VENUS expressing cells from mock (black) and CHIKV-VENKL-infected cell populations (VENUS^+^, red; VENUS^-^, grey). (**E**) Quantification of surface expression and frequency of H2-K^b^ and H2-D^b^ from mock and CHIKV-VENKL-infected EVA cells. Data are pooled (n = 12) (**B**), or representative (n = 9-12) (**C,E**) from 3-4 independent experiments. P values were determined by unpaired student’s t-test (**B**) or one-way ANOVA with Tukey’s test for multiple comparisons (**C,D**). *, *P*<0.05; **, *P*<0.01; ***, *P*<0.001; ****, *P*<0.0001.

After 24 hours (h) in culture, EVA cells were assessed for cellular composition, the capacity to support CHIKV infection, and MHC-I cell surface expression by flow cytometry. More than 95% of EVA cells were CD45^-^ (**Fig 1B and Fig S1A**) and within that population most cells (99%) expressed the fibroblast marker CD29 (**Fig 1C and Fig S1A**), a marker of ankle tissue fibroblasts shown previously to support CHIKV infection *in vivo* (47). In addition, 75-80% of the CD45^-^ cells expressed H2-K^b^ and H2-D^b^ MHC-I molecules on the cell surface (**Fig 1C-D**). To evaluate their capacity to support CHIKV infection, EVA cells were mock-inoculated or inoculated with 10^6^ PFU (plaque forming units) of CHIKV-VENKL and at 24 h post-inoculation (hpi) cells were evaluated for VENUS expression as an indicator of productive CHIKV infection. Consistent with prior studies (47), most CHIKV-infected cells were CD45^-^ and expressed CD29 on their surface (**Fig 1B and Fig S1A**). Next, we assessed surface levels of MHC-I on CHIKV-infected VENUS^+^ cells and control cells. VENUS^+^CD45^-^ EVA cells showed marked reduction of H2-D^b^ and H2-K^b^ cell surface expression compared with VENUS^-^CD45^-^ EVA cells exposed to CHIKV, but not mock-infected CD45^-^ EVA cells, as measured by geometric mean fluorescence intensity (gMFI) (**Fig 1D-E**). The percentage of VENUS^+^CD45^-^ EVA cells that displayed cell surface H2-K^b^ and H2-D^b^ was reduced compared with mock-infected CD45^-^ EVA cells (**Fig 1D-E**). In contrast, the percentage of VENUS^+^CD45^-^ EVA cells that expressed cell surface CD29 was unchanged compared with control cells (**Fig 1B**). To assess the extent to which VENUS expression correlated with cell surface levels of MHC-I, cells were stratified into gates based on the intensity of VENUS expression (**Fig S1B**) and then assessed for H2-K^b^ and H2-D^b^ expression. As VENUS expression increased, H2-Kb and H2-Db expression decreased (**Fig S1C**), suggesting cells with more CHIKV infection express less cell surface MHC-I. Collectively, these data suggest that CHIKV infection suppresses cell surface MHC-I expression but not expression of other cell surface proteins.

### Determinants in nsP2 promote impairment of MHC-I antigen presentation by CHIKV- infected cells

CHIKV nsP2 interferes with numerous cellular functions, including host cell transcription by promoting degradation of RPB1 (15), type 1 IFN responses by nuclear export of STAT1 (16) and inhibition of Jak/STAT phosphorylation (17, 48), and the unfolded protein response (UPR) (18), all of which can be important for MHC-I antigen presentation (49, 50, 51, 52). Given this, we hypothesized that CHIKV nsP2 impairs MHC-I antigen presentation in CHIKV-infected cells. In prior studies, Akhrymuk et al. demonstrated that mutations in the methyltransferase-like domain of CHIKV nsP2 at _ATL_674-676_QMS_ (QMS) abrogate nsP2-mediated inhibition of cellular transcription and type I IFN responses, as well as the cytopathic effects induced by CHIKV infection, without compromising CHIKV replication *in vitro* (53). We therefore incorporated the QMS mutations into CHIKV-VENKL (CHIKV^QMS^-VENKL) (**Fig 2A**) and confirmed that murine fibroblasts supported comparable viral replication of CHIKV-VENKL and CHIKV^QMS^-VENKL, particularly in the absence of type I IFN signaling (**Fig 2B**). EVA cells were inoculated with 10^6^ PFU of CHIKV-VENKL or CHIKV^QMS^-VENKL and at 24 hpi cells were evaluated by flow cytometry for VENUS and cell surface H2-K^b^ (**Fig 2C and Fig S2**). In addition, we quantified cell surface SIINFEKL-loaded H2-K^b^ (H2-K^b^-SIINFEKL) using the 25D1.15 T cell receptor mimic monoclonal antibody (54). CHIKV-VENKL and CHIKV^QMS^-VENKL both infected CD45^-^ EVA cells, albeit with modestly different efficiencies (**Fig 2D and Fig S2**). The surface level of CD29 was similar on cells infected with either CHIKV-VENKL or CHIKV^QMS^-VENKL (**Fig 2E and Fig S2**). In contrast, the surface level of H2-K^b^ on cells infected with CHIKV-VENKL was reduced compared with cells infected with CHIKV^QMS^-VENKL (**Fig 2F**). Moreover, the percentage of VENUS^+^ cells that were double positive for both H2-K^b^-SIINFEKL (i.e., viral peptide-loaded MHC-I) and H2-K^b^ was significantly lower among cells infected with CHIKV- VENKL compared with CHIKV^QMS^-VENKL, and these cells showed markedly lower pMHC-I presentation efficiency (**Fig 2G-H**). These data indicate that MHC-I presentation of viral peptides is less efficient in CHIKV-infected cells and that this can be improved by mutations in nsP2.

**Figure 2.**
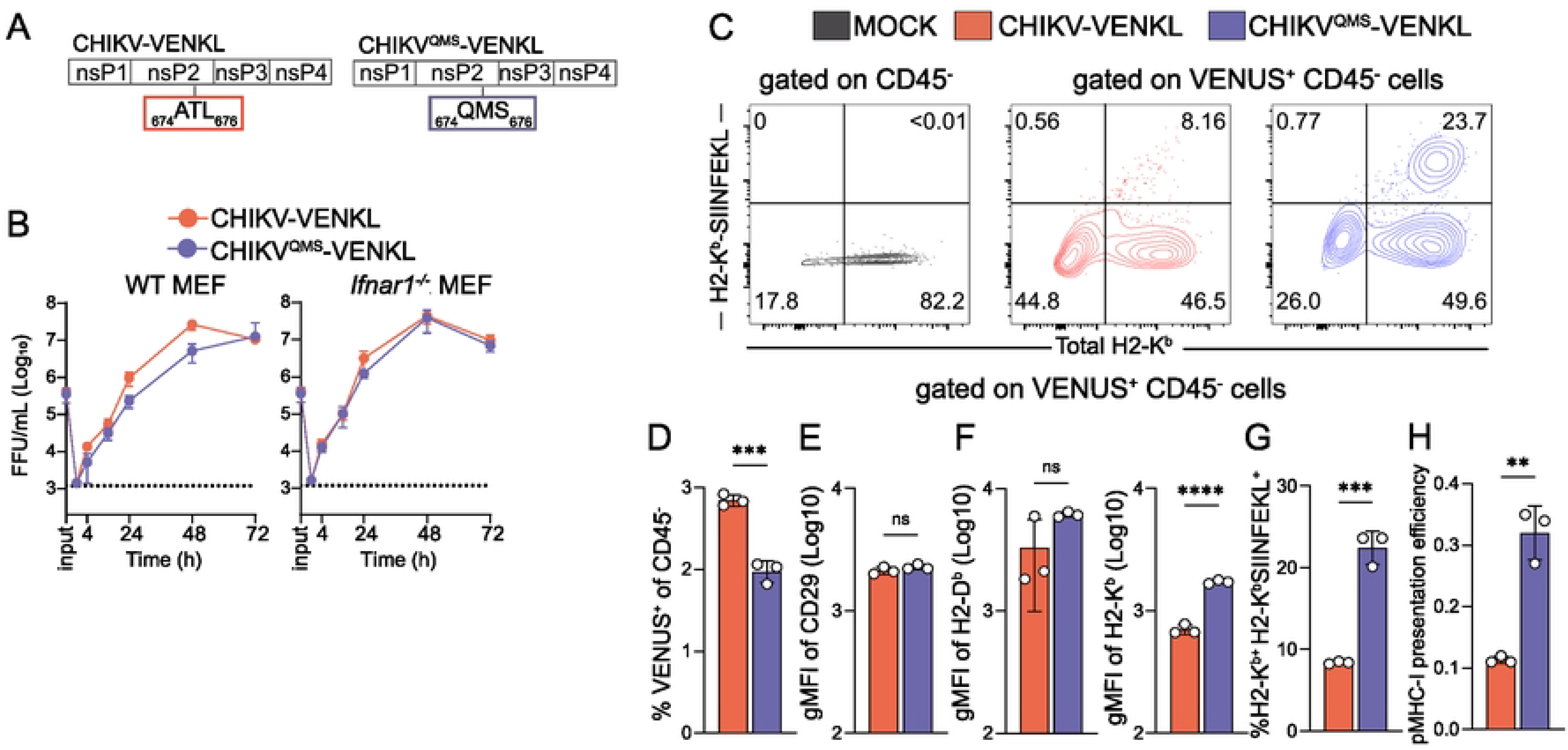
Determinants in nsP2 promote impairment of MHC-I antigen presentation by CHIKV-infected cells. (**A**) Schematic depicting the location of nsP2 V-loop mutations (QMS; blue) compared with wildtype nsP2 (ATL; red). (**B**) WT and *Ifnar1*^-/-^ murine embryonic fibroblasts (MEFs) were inoculated with CHIKV-VENKL or CHIKV^QMS^-VENKL at an MOI of 0.1 FFU/cell. At 0 (input), 1, 24, 48, and 72 hpi, the amount of infectious virus present in culture supernatants was quantified by focus formation assay. (**C-H**) EVA cells were mock-inoculated or inoculated with CHIKV-VENKL or CHIKV^QMS^-VENKL. At 24 hpi, cells were assessed for cell surface expression of H2-K^b^, and SIINFEKL-loaded H2-K^b^ (H2-K^b^-SIINFEKL) by flow cytometry. (**C**) Representative flow cytometry plots depicting H2-K^b^-SIINFEKL and H2-K^b^ cell surface expression on live, CD45^-^ and live, CD45^-^, VENUS^+^ cells from mock-, CHIKV-VENKL-, and CHIKV^QMS^-VENKL-inoculated EVA cells. (**D**) Percentage of VENUS^+^ cells among live, CD45^-^, EVA cells. (**E**) CD29 cell surface expression on live, CD45^-^, VENUS^+^ EVA cells. (**F**) Quantification of H2-D^b^ and H2-K^b^ surface expression on live, CD45^-^, VENUS^+^ EVA cells. (**G**) Percentage of double-positive (H2-K^b+^H2-K^b^-SIINFEKL^+^) cells on live, CD45^-^, VENUS^+^ EVA cells. (**H**) pMHC-I presentation efficiency on live, CD45^-^, VENUS^+^ EVA cells. Data are representative of 4 experiments (n = 12). P values were determined by unpaired student’s t-test (**D**). **, *P*<0.01; ***, *P*<0.001.

To further assess whether CHIKV infection interferes with functional MHC-I antigen presentation, and validate our direct assessment of cell surface H2-Kb-SIINFEKL detected by the 25D1.16 antibody, we developed a co-culture system utilizing highly specific and sensitive transgenic OT-I CD8^+^ T cells, which upregulate cell surface expression of CD69 and CD25 upon recognition of and activation by H2-K^b^-SIINFEKL complexes (55). To determine if OT-I CD8^+^ T cells are activated by CHIKV-infected cells, EVA cells were inoculated with mock inoculum or 10^6^ PFU of CHIKV-VENKL or CHIKV^QMS^-VENKL. At 24 hpi, all groups of cells were washed and co-cultured for 6 h with a 1:1 mixture of 10^6^ purified OT-I CD8^+^ T cells and 10^6^ bystander CD8^+^ T cells with >95% purity and minimal background activation prior to addition (**Fig S3A-S3C**). Consistent with the low level of cell surface H2-K^b^-SIINFEKL detected on CHIKV-VENKL-infected EVA cells (**Fig 2C and 2G**), OT-I CD8^+^ T cells co-cultured with CHIKV-VENKL-infected EVA cells showed significantly lower fractions of CD69 and CD25 double positive cells (**Fig 3A and 3B**), and surface expression of CD69 (**Fig 3C**) and CD25 (**Fig 3D**) than CHIKV^QMS^-VENKL- infected EVA cells. OT-I CD8^+^ T cells co-cultured with CHIKV-VENKL infected EVA cells had increased cell surface CD69 expression compared with bystander CD8^+^ T cells (**Fig 3A and 3C**), however, few of these cells expressed CD25 (**Fig 3A**) indicating that their activation was not specific to TCR ligation. These data suggest that CHIKV-infected cells are inefficiently recognized by antigen-specific CD8^+^ T cells and that this evasion is promoted by determinants in nsP2. In addition, these findings demonstrate that the lower signal detected by the H2-K^b^-SIINFEKL antibody on CHIKV-VENKL-infected cells functionally impacts antigen presentation to CD8^+^ T cells.

**Figure 3.**
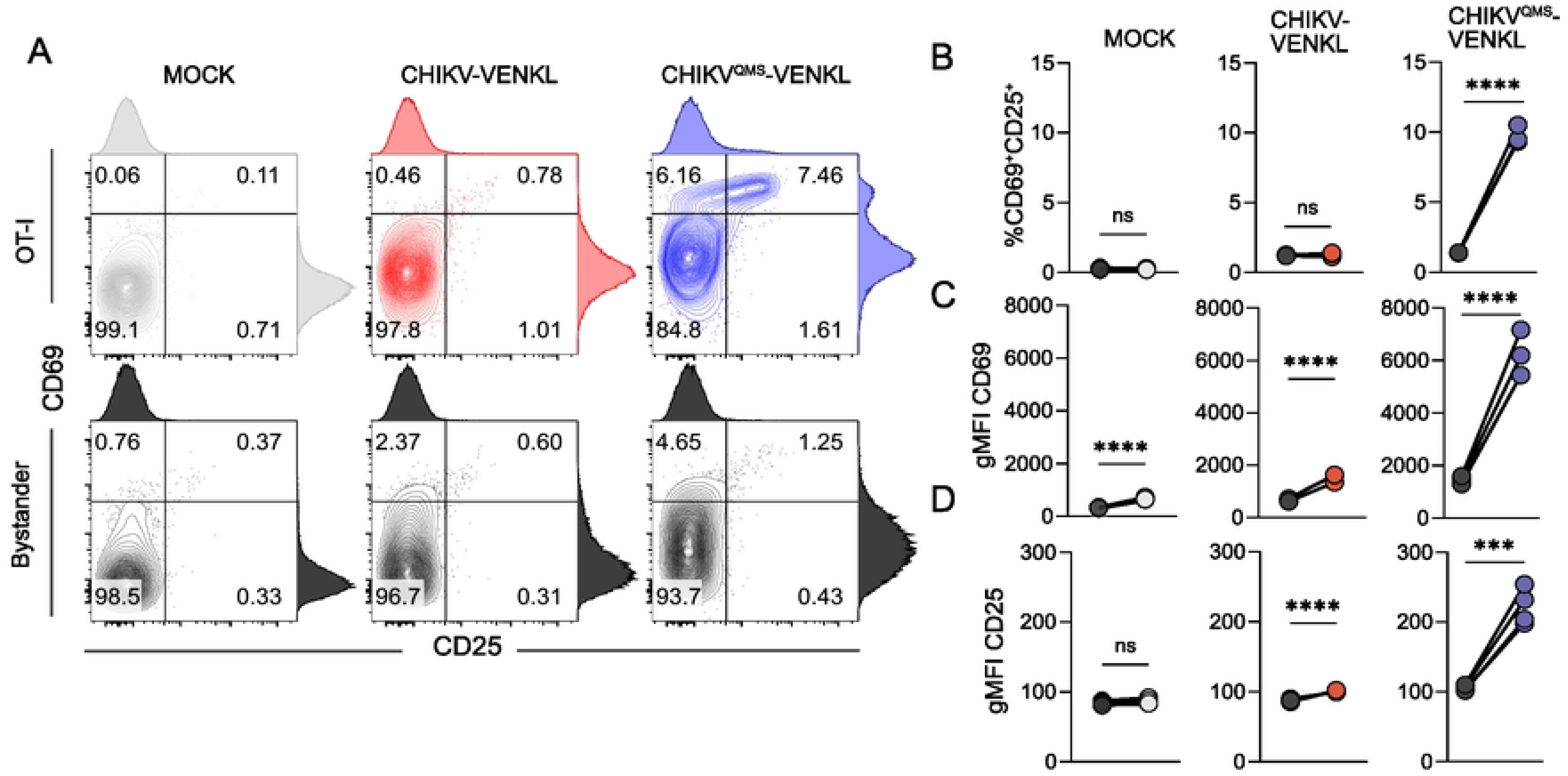
WT CHIKV-infected cells inefficiently activate CD8^+^ T cells *in vitro*. (**A-D**) EVA cells were mock-inoculated or inoculated with 10^6^ PFU of CHIKV-VENKL or CHIKV^QMS^- VENKL. At 24 hpi, EVA cells were cocultured with a 1:1 mixture of 10^6^ bystander CD8^+^ T cells and 10^6^ OT-I SIINKEKL-specific CD8^+^ T cells for 6 h. Bystander CD8^+^ T cells were distinguished from OT-I CD8^+^ T cells by gating on CD8^+^, KbSIINFEKL tetramer positive or negative cells, and both populations were assessed for the dual expression of CD69 and CD25 by flow cytometry. (**A**) Representative flow cytometry plots. (**B**) Quantification of CD25^+^CD69^+^ double positive cell frequencies for bystander (black) and OT-I (colored) CD8^+^ T cells. (**C**) Cell surface expression of CD69 (gMFI). (**D**) Cell surface expression of CD25 (gMFI). Data are representative of two independent experiments (n = 6). P values were determined by paired student’s t-test. ****, *P*<0.0001.

### MHC-I antigen presentation is inefficient in CHIKV-infected cells *in vivo*

To evaluate whether CHIKV-infected cells display poor MHC-I antigen presentation *in vivo*, WT C57BL/6 mice were inoculated in the left-rear footpad with 10^3^ PFU of either CHIKV-VENKL or CHIKV^QMS^-VENKL and ankle-associated cells were evaluated by flow cytometry for VENUS, as an indicator of infection (**Fig 4A and Fig S4**), and cell surface CD45, CD29, CD44, H2-K^b^ and H2-K^b^-SIINFEKL expression on day 1, 2, and 5 post inoculation. Like EVA cells infected *ex vivo*, 85-95% of VENUS^+^ cells in both CHIKV-VENKL and CHIKV^QMS^-VENKL infected mice were CD45^-^CD29^+^ (**Fig 4B and Fig S4**), indicating that most cells infected by either virus are joint tissue-associated fibroblasts. However, fewer infected cells were detected at both 1 and 2 dpi in mice inoculated with CHIKV^QMS^-VENKL (**Fig 4A and Fig S4**), suggesting replication of this virus is restricted. At 2 and 5 dpi, CHIKV-VENKL-infected cells had lower cell surface expression of H2-K^b^, as determined by the ratio of H2-K^b^ gMFI on VENUS^+^ cells to VENUS^-^ cells, compared with CHIKV^QMS^-VENKL-infected cells (**Fig 4C**). While CD44 (**Fig 4D**) and CD29 (**Fig 4E**) expression on CHIKV-VENKL- and CHIKV^QMS^-VENKL-infected cells eventually differ by 5 dpi, at 1 and 2 dpi cell surface expression of both molecules was comparable. In comparison with CHIKV^QMS^-VENKL-infected cells, a lower percentage of CHIKV-VENKL-infected cells displayed H2-K^b^-SIINFEKL on their surface across all time points examined, resulting in reduced pMHC-I presentation efficiency (**Fig 4F-H**).

**Figure 4.**
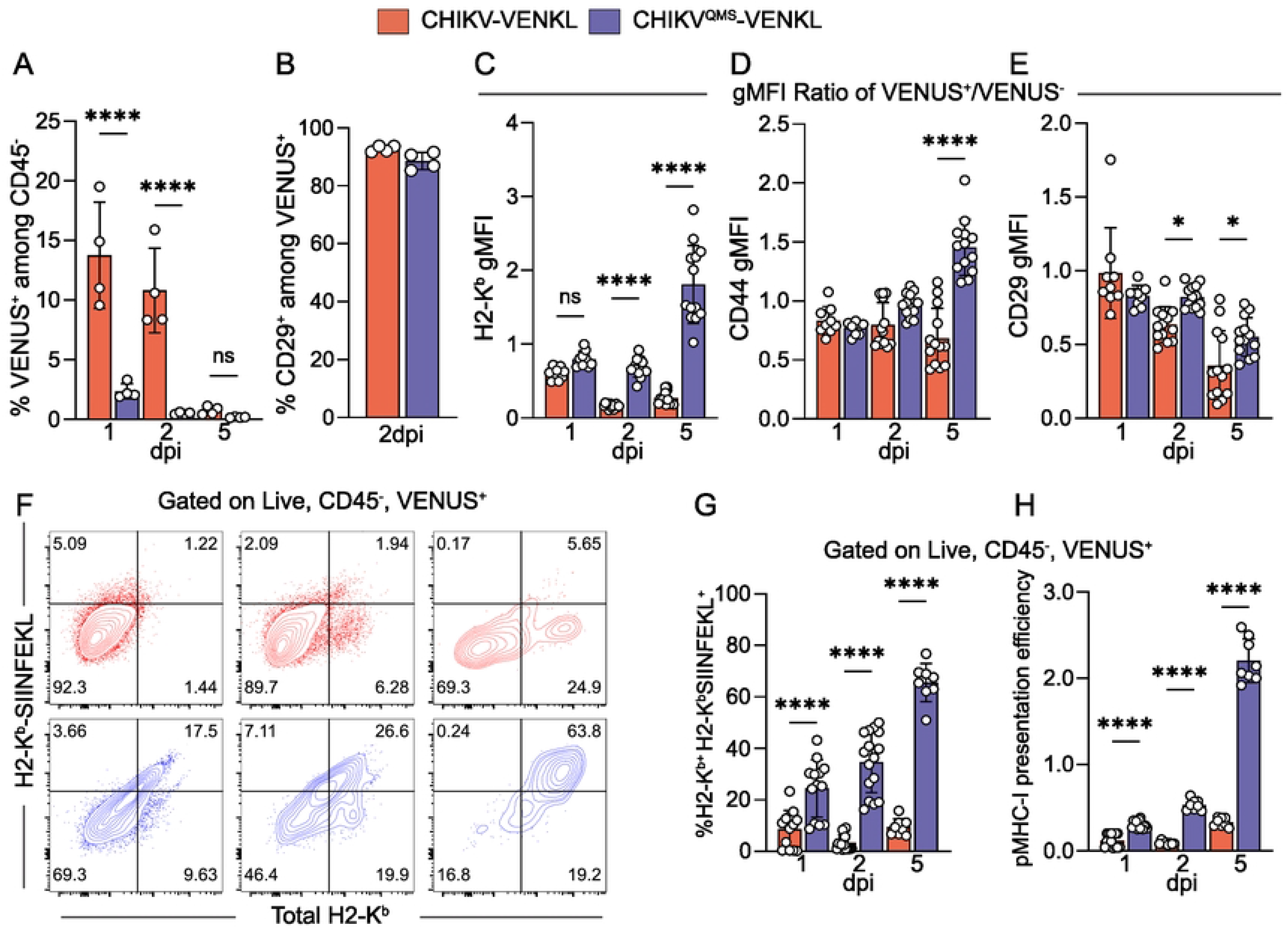
CHIKV-infected cells inefficiently display MHC-I antigen *in vivo*. (**A-H**) WT C57BL/6 mice were inoculated with 10^3^ PFU of CHIKV-VENKL (red) or CHIKV^QMS^-VENKL (blue). At 1, 2, and 5 dpi, total cells were isolated from the ipsilateral foot and ankle and evaluated by flow cytometry. (**A**) Frequency of VENUS^+^ cells among live, CD45^-^ cells. (**B**) Frequency of CD29^+^ cells among live, CD45^-^, VENUS^+^ cells at 2 dpi. (**C-E**) Cell surface expression of H2-K^b^, CD44, and CD29 among live, CD45^-^, VENUS^+^ cells normalized to VENUS^-^ cells. (**F**) Representative flow cytometry plots of total H2-K^b^ and H2-K^b^-SIINFEKL staining. (**G**) Quantification of the frequency of double-positive (H2-K^b+^H2-K^b^-SIINFEKL^+^) cells among live, CD45^-^, VENUS^+^ cells. (**H**) pMHC-I presentation efficiency among live, CD45^-^, VENUS^+^ cells. Data are representative of 3 experiments (n = 12) (**A-B**) or pooled from 3-4 independent experiments (C-H) (n = 8-13). P values were determined by one-way ANOVA with Tukey’s multiple comparison test (**B**). *, *P*<0.05; ****, *P*<0.0001.

### Cell surface, but not cell-associated, MHC-I expression is reduced in CHIKV-infected cells

CHIKV nsP2 impairs cellular transcription *in vitro* (56, 57) which could explain the poor pMHC-I presentation efficiency by CHIKV-infected cells and the increase in pMHC-I presentation by cells infected with CHIKV encoding the QMS mutations in nsP2. To investigate whether the suppression of cell surface MHC-I expression in CHIKV-infected cells is due to a reduction in total cell-associated MHC-I, we developed a flow cytometry-based intracellular staining protocol for H2-K^b^ to allow for concurrent assessment of cell surface and cell-associated MHC-I on single EVA cells (**Fig 5A**). The intracellular quantity of H2-K^b^ and percentage of cells expressing intracellular H2-K^b^ was similar for CHIKV-VENKL- and CHIKV^QMS^-VENKL-infected cells. In contrast, fewer CHIKV-VENKL-infected cells displayed H2-K^b^ on their surface, and those that did had reduced expression levels (**Fig 5B-C**). These data suggest that CHIKV infection does not downregulate expression of H2-Kb by infected cells, but instead disrupts a step of the MHC-I antigen presentation pathway required for the display of mature pMHC-I on the cell surface.

**Figure 5.**
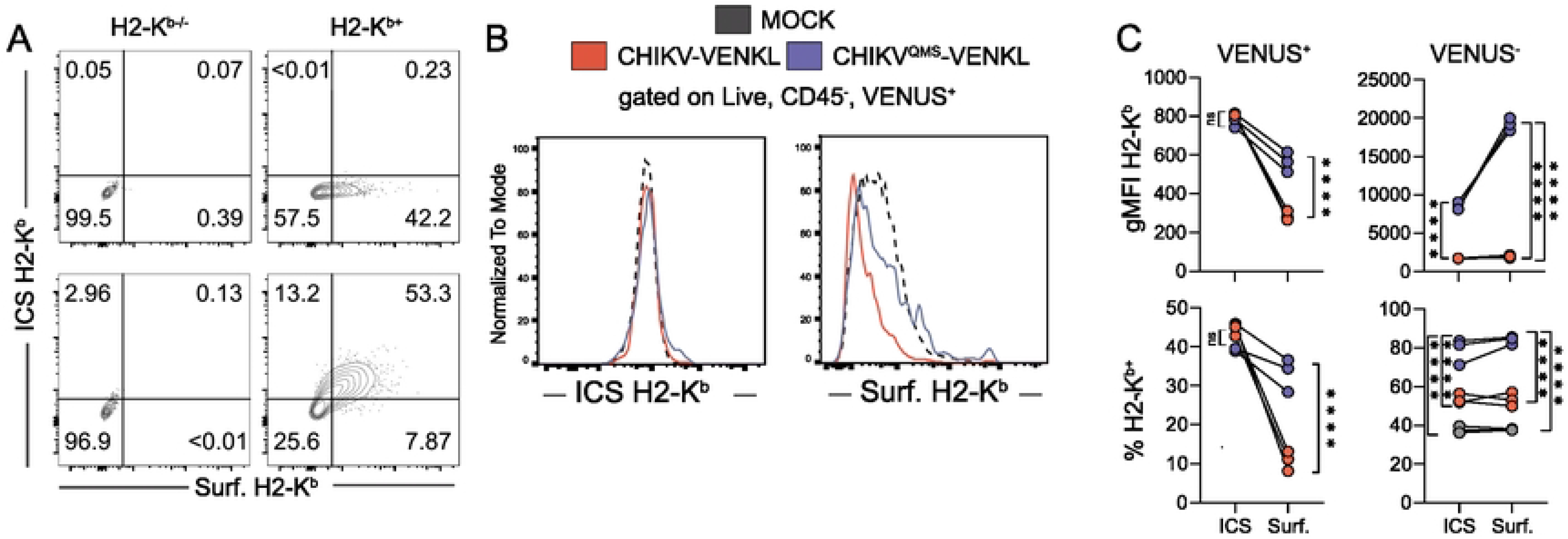
Cell surface, but not cell-associated, MHC-I expression is reduced in CHIKV-infected cells. (**A**) WT murine embryonic fibroblasts (MEFs) stably overexpressing H2- K^b^ (H2-K^b+^) and H2-K^b-/-^ MEFs were analyzed for intracellular and cell surface expression of H2-K^b^ by flow cytometry (ICS= intracellular stain, Surf. = surface stain). (**B-C**) EVA cells were mock-inoculated or inoculated with 10^6^ PFU of CHIKV-VENKL or CHIKV^QMS^-VENKL. At 24 hpi, cells were analyzed for intracellular (ICS) and cell surface (Surf.) expression of H2-K^b^ by flow cytometry. (**B**) Representative flow cytometry plots and (**C**) quantification of the magnitude (gMFI) and frequency (%) of intracellular (ICS) and cell surface (Surf.) H2-K^b^ expression on CHIKV-infected cells (VENUS^+^) and VENUS^-^ cells. Data are representative of 2 independent experiments (n = 6). P values were determined by unpaired student’s t-test. ****, *P*<0.0001.

### Bypassing peptide processing and ER transport restores MHC-I antigen presentation by CHIKV-infected cells

Given our findings that CHIKV-infected cells inefficiently present pMHC-I, and that intracellular expression of H2-K^b^ is normal during CHIKV-VENKL infection, we sought to determine the step(s) at which MHC-I maturation is disrupted in CHIKV-infected cells. Mature MHC-I complexes are composed of a membrane anchored heavy chain (e.g., H2-K^b^), in association with beta-2-microgobulin (β_2_m), bound to an 8-10 amino acid peptide. β_2_m and peptide of proper length are required for the heavy chain of MHC-I to traffic to the cell surface (58), and the absence of an appropriate peptide destabilizes MHC-I during transit and at the cell surface (59, 60). Prior studies found that the covalent linkage of an antigenic peptide to β_2_m via a flexible linker allows for assembly of mature, peptide-loaded MHC-I complexes independent of peptide processing, transporter associated with antigen processing (TAP)-mediated peptide translocation into the ER lumen, and the peptide loading complex (PLC) (61, 62, 63). To evaluate if CHIKV-infected cells are deficient for these early steps in MHC-I antigen presentation, the VENKL coding sequence in CHIKV-VENKL was replaced with SIINFEKL-GSSGSSGS-β_2_m (CHIKV-SIINFEKLβ_2_m) (**Fig 6A**). EVA cells were mock-inoculated or inoculated with CHIKV-VENKL, CHIKV^QMS^-VENKL, or CHIKV-SIINFEKLβ_2_m. At 24 hpi, CHIKV-infected cells were stained with an anti-E2 monoclonal antibody (CHK-11), which identified CHIKV-infected cells as efficiently as detection of virus-encoded VENUS (**Fig S5**), and virus-infected cells were evaluated for cell surface H2-K^b^ and H2-K^b^-SIINFEKL expression. In comparison with CHIKV-VENKL-infected EVA cells, an increased percentage of CHIKV-SIINFEKLβ_2_m-infected cells displayed H2-K^b^-SIINFEKL on the cell surface, resulting in improved pMHC-I presentation efficiency that was similar to that observed in CHIKV^QMS^-VENKL- infected cells (**Fig 6B and 6C**). Likewise, the inoculation of mice with CHIKV-SIINFEKLβ_2_m elicited increased pMHC-I presentation efficiency and an increased percentage of antigen presenting CHIKV-infected cells compared to CHIKV-VENKL at 2 dpi, even above that seen by cells infected with CHIKV^QMS^-VENKL (**Fig 6D and 6E**). These findings suggest CHIKV infection disables MHC-I antigen presentation at a step prior to β_2_m-peptide stabilization of the heavy chain within the ER.

**Figure 6.**
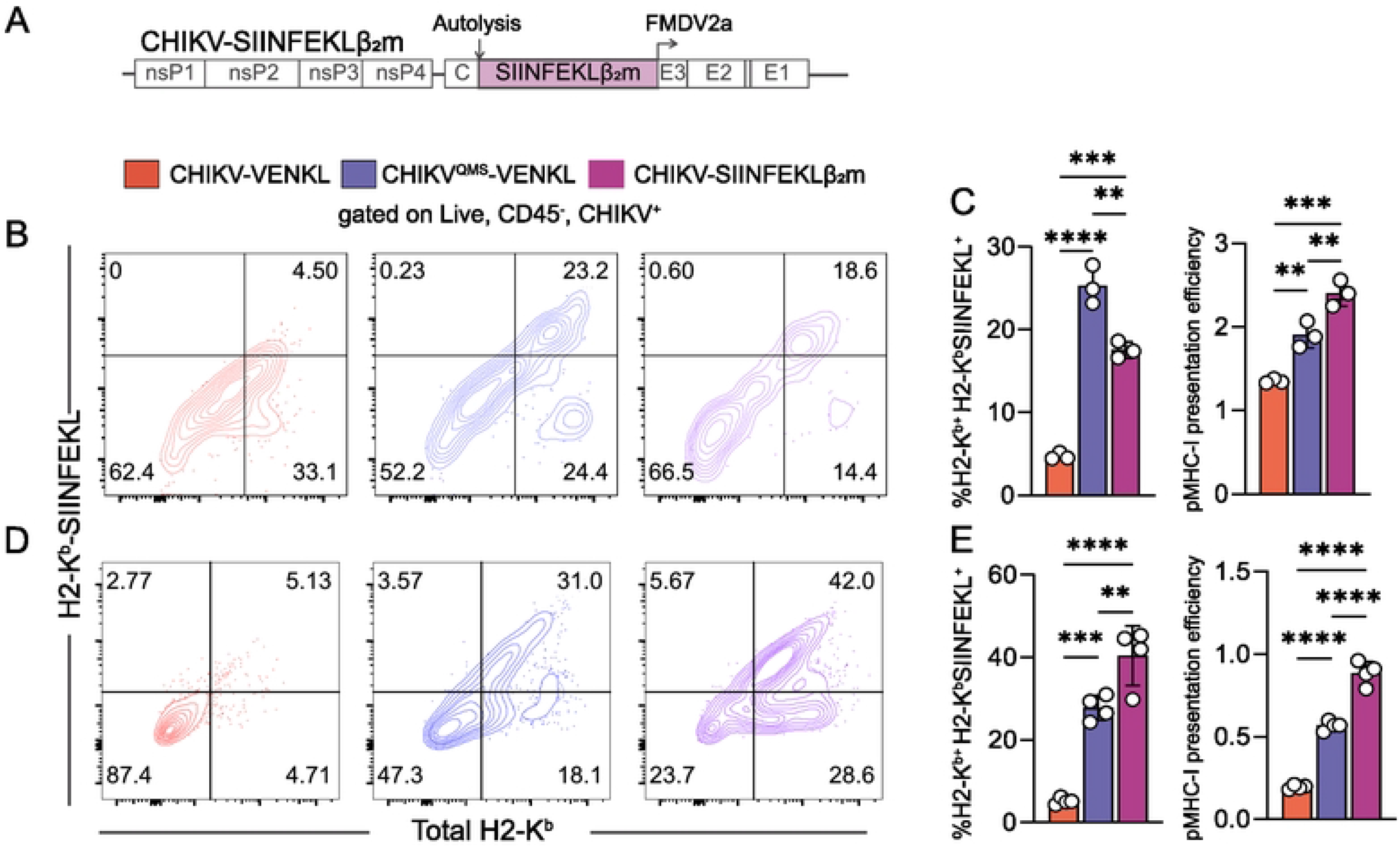
Bypassing peptide processing and ER transport restores MHC-I antigen presentation by CHIKV-infected cells. (**A**) Schematic of the recombinant CHIKV-SIINFEKLβ_2_m viral genome. The coding sequence for the β_2_mSIINFEKL chimeric protein was inserted into the CHIKV genome in-frame in the viral structural ORF similar to the VENKL coding sequence. (**B and C**) EVA cells were mock-inoculated or inoculated with CHIKV-VENKL, CHIKV^QMS^-VENKL, or CHIKV-SIINFEKLβ_2_m. At 24 hpi, live, CD45^-^, E2 (CHIKV)^+^ cells were assessed for cell surface expression of H2-K^b^ and SIINFEKL-loaded H2-K^b^ (H2-K^b^-SIINFEKL) by flow cytometry. (**B**) Representative flow cytometry plots depicting the frequency of double-positive (H2-K^b+^H2-K^b^-SIINFEKL^+^) cells among live, CD45^-^, CHIKV^+^ cells. (**C**) Quantification of the frequency of double-positive (H2-K^b+^H2-K^b^-SIINFEKL^+^) cells (left) and pMHC-I presentation efficiency (right). (**D and E**) At 24 hpi, ankle cells from C57BL/6 mice mock-inoculated or inoculated with 10^6^ PFU CHIKV-VENKL, CHIKV^QMS^-VENKL, or CHIKV-SIINFEKLβ_2_m were assessed by flow cytometry. (**D**) Representative flow cytometry plots depicting the frequency of double-positive (H2-K^b+^H2-K^b^-SIINFEKL^+^) cells among live, CD45^-^, CHIKV^+^ cells. (**E**) Quantification of the frequency of double-positive (H2-K^b+^H2-K^b^-SIINFEKL^+^) cells (left) and pMHC-I presentation efficiency (right) among live, CD45^-^, CHIKV^+^ cells. Data are representative of 2 independent experiments (n = 8). P values were determined by one-way ANOVA with Tukey’s multiple comparison test. **, *P*<0.01; ***, *P*<0.001; ****, *P*<0.0001.

To further demonstrate that presentation of viral antigen by MHC-I can occur more efficiently in CHIKV-infected cells when the peptide is linked to β_2_m, EVA cells were inoculated with mock inoculum or 10^6^ PFU of CHIKV-VENKL, CHIKV^QMS^-VENKL, or CHIKV-SIINFEKLβ_2_m. At 24 hpi, all groups of cells were washed and co-cultured for 6 h with a 1:1 mixture of 10^6^ purified OT-I CD8^+^ T cells and 10^6^ bystander CD8^+^ T cells. Like OT-I CD8^+^ T cells co-cultured with CHIKV^QMS^-VENKL-infected EVA cells, OT-I CD8^+^ T cells co-cultured with CHIKV-SIINFEKLβ_2_m-infected cells displayed increased expression of activation markers CD69 and CD25 compared with bystander levels (**Fig 7A-B**) and OT-I CD8^+^ T cells co-cultured with CHIKV-VENKL-infected EVA cells. Furthermore, these OT-I CD8^+^ T cells expressed elevated levels of IRF4, a transcription factor activated in response to TCR ligation strength (64), suggesting that specific engagement of CHIKV-infected cells with OT-I CD8^+^ T cells is restored when the virus-encoded peptide is directly linked with β_2_m. Notably, OT-I CD8^+^ T cells cocultured with CHIKV-SIINFEKLβ_2_m-, but not CHIKV-VENKL-infected EVA cells, showed a marked loss of CD8a expression (**Fig 7B**), which has been observed on activated CD8^+^ T cells (65), further suggesting that antigen-specific CD8^+^ T cells poorly engage cells infected with CHIKV-VENKL.

**Figure 7.**
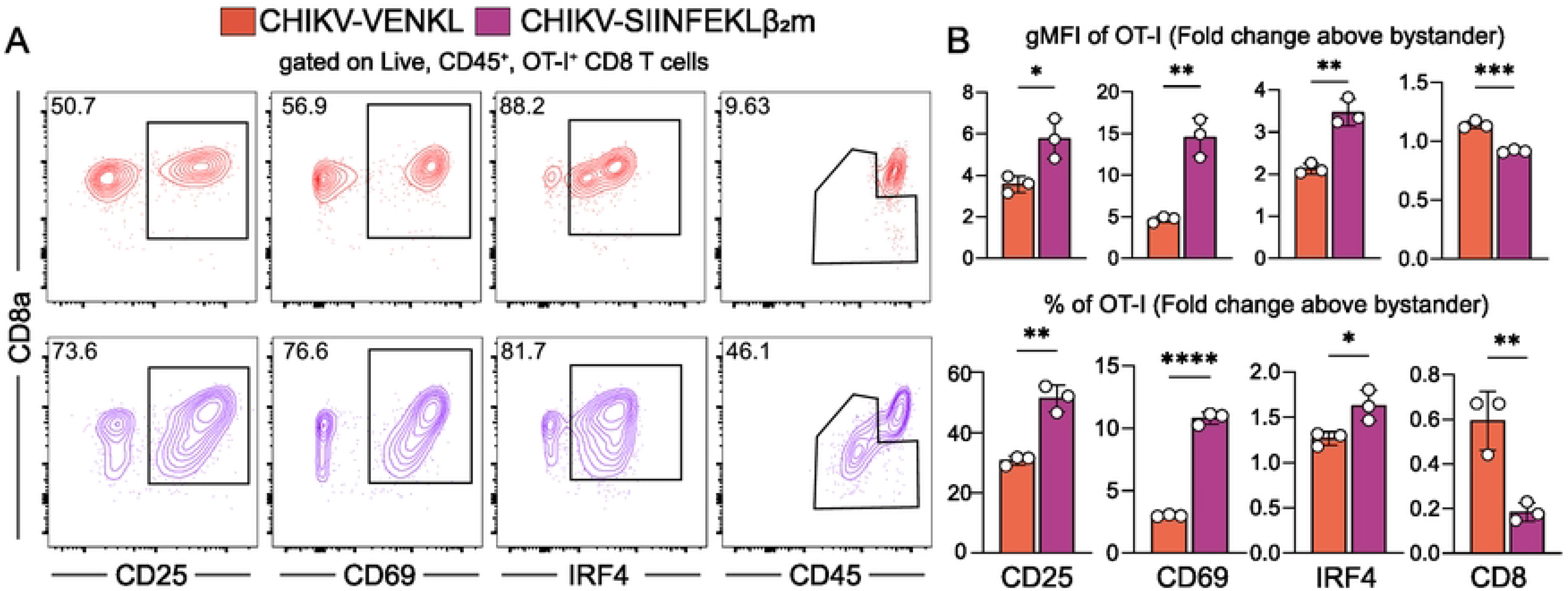
CD8^+^ T cells are activated by CHIKV-infected cells when peptide processing and ER transport are bypassed. (**A and B**) EVA cells were inoculated with 10^6^ PFU CHIKV-VENKL or CHIKV-SIINFEKLβ_2_m. At 24 hpi, EVA cells were cocultured with a 1:1 mixture of 10^6^ bystander and 10^6^ OT-I SIINKEKL-specific CD8^+^ T cells for 6 h. Bystander CD8^+^ T cells were distinguished from OT-I CD8^+^ T cells by gating on CD8^+^, KbSIINFEKL tetramer positive or negative cells, and both populations were assessed for the expression of CD69, CD25, IRF4 and CD8 by flow cytometry. (**A**) Representative flow cytometry plots. (**B**) Quantification of CD25, CD69, IRF4 (intracellular), and CD8 cell surface expression (upper graphs) and frequency (lower graphs) for OT-I cells above bystander CD8^+^ T cells. Data are representative of two independent experiments (n = 6). P-values were determined by unpaired student’s t-test. *, *P*<0.05; **, *P*<0.01; ***, *P*<0.001; ****, *P*<0.0001.

### CHIKV nsP2 impairs antigen presentation efficiency

Our data shows that mutations in the methyltransferase-like domain of nsP2 increased MHC-I viral antigen presentation by CHIKV-infected cells. Based on these data, we hypothesized that nsP2 expression alone would be sufficient to disrupt pMHC-I presentation efficiency. To test this idea, EVA cells were co-transfected with a CHIKV nsP2 expression vector along with an expression vector encoding VENKL. Control cells were co-transfected with a control vector plus VENKL. At 24 h post-transfection, ∼20-30% of EVA cells were VENUS^+^ (**Fig 8A**). Importantly, expression of both nsP2 and nsP2^QMS^ were detected at similar levels by Western blot (**Fig 8B**). In cells transfected with nsP2, VENUS^+^CD45^-^ EVA cells displayed a reduced percentage of cells double-positive for H2-K^b^ and H2-K^b^-SIINFEKL, and decreased pMHC-I presentation efficiency (**Fig 8C-D**) compared with control cells. In contrast, EVA cells transfected with nsP2^QMS^ and VENKL had a similar frequency of the frequency of double-positive (H2-K^b+^H2-K^b^-SIINFEKL^+^) cells and pMHC-I presentation efficiency as control cells (**Fig 8C-D**). These findings indicate that nsP2 expression alone can impair pMHC-I presentation efficiency similar to CHIKV-infected cells.

**Figure 8.**
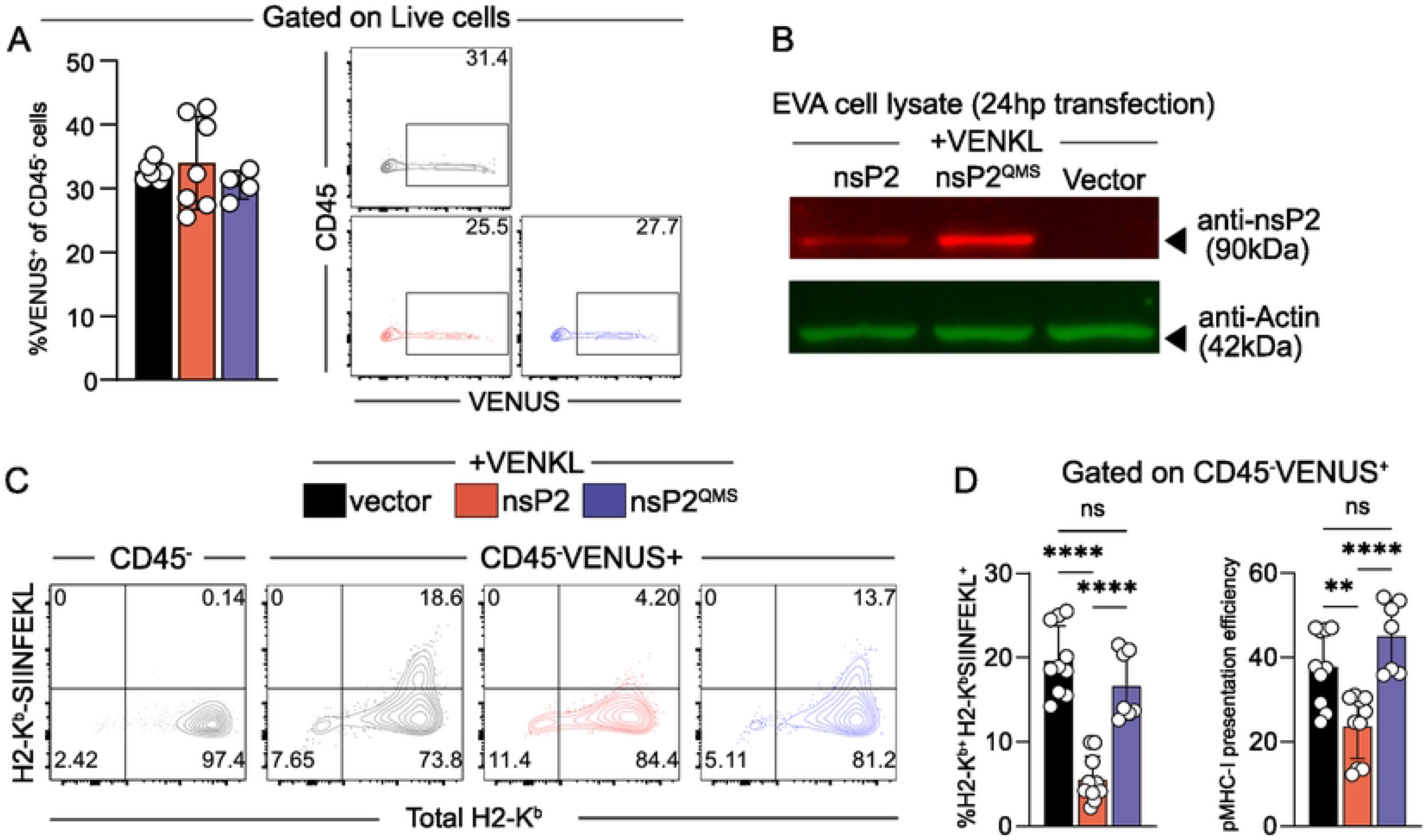
Cells expressing nsP2 inefficiently present antigen and have decreased levels of cell surface MHC-I. (**A-D**) EVA cells were co-transfected with pCMV-nsP2 or an identical control plasmid lacking the methionine start codon for nsP2 (vector) and pCMV-VENKL. At 24 h post-transfection, cells were evaluated by flow cytometry and Western blot. (**A**) VENUS^+^ cells among live, CD45^-^ cells and representative flow cytometry plots. (**B**) Western blot analysis of nsP2 and actin expression within transfected EVA cells (anti-nsP2, red, 90 kDa; anti-actin, green, 42 kDa). (**C**) Representative flow cytometry plots depicting the frequency of double-positive (H2-K^b+^H2-K^b^-SIINFEKL^+^) cells among live, CD45^-^, VENUS^+^ cells. (**D**) Quantification of the frequency of double-positive (H2-K^b+^H2-K^b^-SIINFEKL^+^) cells and pMHC-I presentation efficiency among live, CD45^-^, VENUS^+^ cells. Data are pooled from 3 independent experiments (n = 9). P values were determined by one-way ANOVA with Tukey’s multiple comparisons test. **, *P*<0.01; ****, *P*<0.0001.

## DISCUSSION

Replication of arthritogenic alphaviruses in musculoskeletal tissues of mice, nonhuman primates, and humans leads to the recruitment and infiltration of inflammatory cells (66, 67, 68, 69, 70, 71). Some of these cellular infiltrates, such as Ly6C^hi^ monocytes, contribute to control of alphavirus infection (72, 73), whereas others, such as CD4^+^ T cells, mediate joint tissue pathology (74, 75). These tissue infiltrates also include CD8^+^ T cells (41), which are critical for the control and clearance of many viral infections. Remarkably, prior studies found that CD8^+^ T cells have little to no impact on CHIKV or RRV replication in joint-associated tissues (41, 76). However, CD8^+^ T cells in joint-associated tissues of CHIKV-infected mice were capable of lysing adoptively transferred splenocytes exogenously loaded with peptide (41), suggesting that CHIKV-infected cells in joint tissue escape CD8^+^ T cell surveillance. Thus, we investigated MHC-I antigen presentation by CHIKV-infected cells. To do this, we engineered a recombinant CHIKV strain (CHIKV-VENKL) that allowed us to quantify MHC-I surface expression and antigen presentation efficiency of a virus-encoded peptide by infected cells. This virus expresses the VENUS fluorescent protein reporter with the SIINFEKL CD8^+^ T cell epitope embedded within its open reading frame allowing for concurrent determination of infection status and aspects of MHC-I antigen presentation. Because SIINFEKL within the VENUS protein is efficiently liberated by proteasomes and trimmed for MHC-I binding, pMHC-I presentation efficiency (i.e., H2-K^b^-SIINFEKL per source protein) can be calculated. Indeed, we found that joint-associated CHIKV-infected fibroblasts displayed reduced cell surface MHC-I expression and inefficiently presented a virus-encoded peptide in MHC-I both *in vitro* and *in vivo*. Moreover, CHIKV-infected fibroblasts were unable to activate viral antigen-specific CD8^+^ T cells, further indicating that MHC-I antigen presentation by CHIKV-infected cells is disrupted.

Viruses have evolved numerous strategies to evade the MHC-I antigen presentation pathway, including disruption of MHC-I machinery by degradation or suppression of transcription (34, 35, 77, 78, 79), highlighting the importance of CD8^+^ T cells in the elimination of virus-infected cells and control of virus infection. Infection of cells with CHIKV and other alphaviruses can inhibit host cell transcription (80). Mechanistically, for CHIKV and many other alphaviruses, inhibition of transcription is mediated by translocation of the viral nsP2 protein into the nucleus where it promotes polyubiquitination and proteasomal degradation of RPB1 (15), the catalytic subunit of cellular DNA-dependent RNA polymerase II. Recently, CHIKV strains that lack the capacity to induce degradation of RPB1 but retain high levels of replication were developed – these strains encode specific mutations in the V-loop of the methyltransferase-like domain of nsP2 (53). We introduced these mutations into CHIKV-VENKL and found that cells infected with this virus retained cell surface expression of MHC-I, displayed more efficient MHC-I antigen presentation, and activated antigen-specific CD8^+^ T cells. Moreover, we found that transfection of primary murine ankle tissue fibroblasts with WT CHIKV nsP2, but not CHIKV nsP2 with mutations in the V-loop, was sufficient to diminish MHC-I surface expression and pMHC-I presentation efficiency. Thus, our studies identified a unique role for nsP2 in immune evasion.

Given that CHIKV infection and nsP2 expression can inhibit host cell transcription, and mutations in nsP2 abrogated disruption of MHC-I antigen presentation in CHIKV-infected and nsP2-expressing cells, we developed a flow cytometry-based assay for quantifying cell-associated and cell surface MHC-I expression on individual cells to determine if CHIKV infection specifically affects cell surface MHC-I expression or induces a more general downregulation of MHC-I. These studies demonstrated that while cell surface expression of MHC-I was decreased in CHIKV-infected cells, intracellular levels of MHC-I were unaffected, indicating that CHIKV infection does not result in a general loss of MHC-I expression due to inhibition of host cell transcription.

Our findings suggest that CHIKV nsP2 interferes with MHC-I antigen presentation by a mechanism that prevents display of mature peptide-loaded MHC-I on the cell surface (**Fig 9**). Prior studies by other groups found that when an MHC-I epitope is covalently tethered to β_2_m, this epitope is translocated into the ER by the β_2_m signal sequence, thus, the presentation of the tethered epitope does not require TAP-mediated transport (61). In addition, this approach bypasses the need for peptide processing (63). We engineered CHIKV to express SIINFEKL tethered to the N-terminus of β_2_m via a Gly/Ser linker (61) (CHIKV-SIINFEKLβ_2_m) and found that cells infected with this virus efficiently display SIINFEKL-loaded MHC-I on the cell surface. Furthermore, the capacity for viral expression of β_2_m-SIINFEKL to restore peptide-loaded MHC-I on the cell surface and promote activation of antigen-specific CD8^+^ T cells indicates that MHC-I trafficking is not altered by CHIKV-infected cells and further supports the idea that steps prior to MHC-I heavy chain stabilization and trafficking are disrupted. Mechanistically, these findings indicate that CHIKV infection and nsP2 expression disrupt MHC-I antigen presentation by altering peptide generation, peptide transport into the ER, or the peptide loading complex (PLC), and future studies are needed to resolve the exact mechanism of action. It is possible that nsP2 directly interacts with components of the antigen processing pathway to disrupt MHC-I antigen presentation. For example, ubiquinilin 4 (UBQLN4) directly interacts with nsP2 (81) and is known to target mis-localized membrane proteins to the proteasome (82). Given the critical importance of the proteasome for the generation of CD8^+^ T cell epitopes (83), it is possible that the interaction between nsP2 and UBQLN4 leads to a reduction of pMHC-I antigenic epitopes for surveillance of CHIKV-infected cells. Alternatively, in contrast to the MHC-I heavy chain, other components of the MHC-I antigen presentation pathway, such as the TAP transporters or the PLC, may be more sensitive to nsP2-mediated transcriptional inhibition. In summary, we find that CHIKV-infected joint tissue fibroblasts exhibit inefficient MHC-I presentation of virus-encoded peptides. These data provide one explanation for the limited role of CD8^+^ T cells in controlling CHIKV infection.

**Figure 9.**
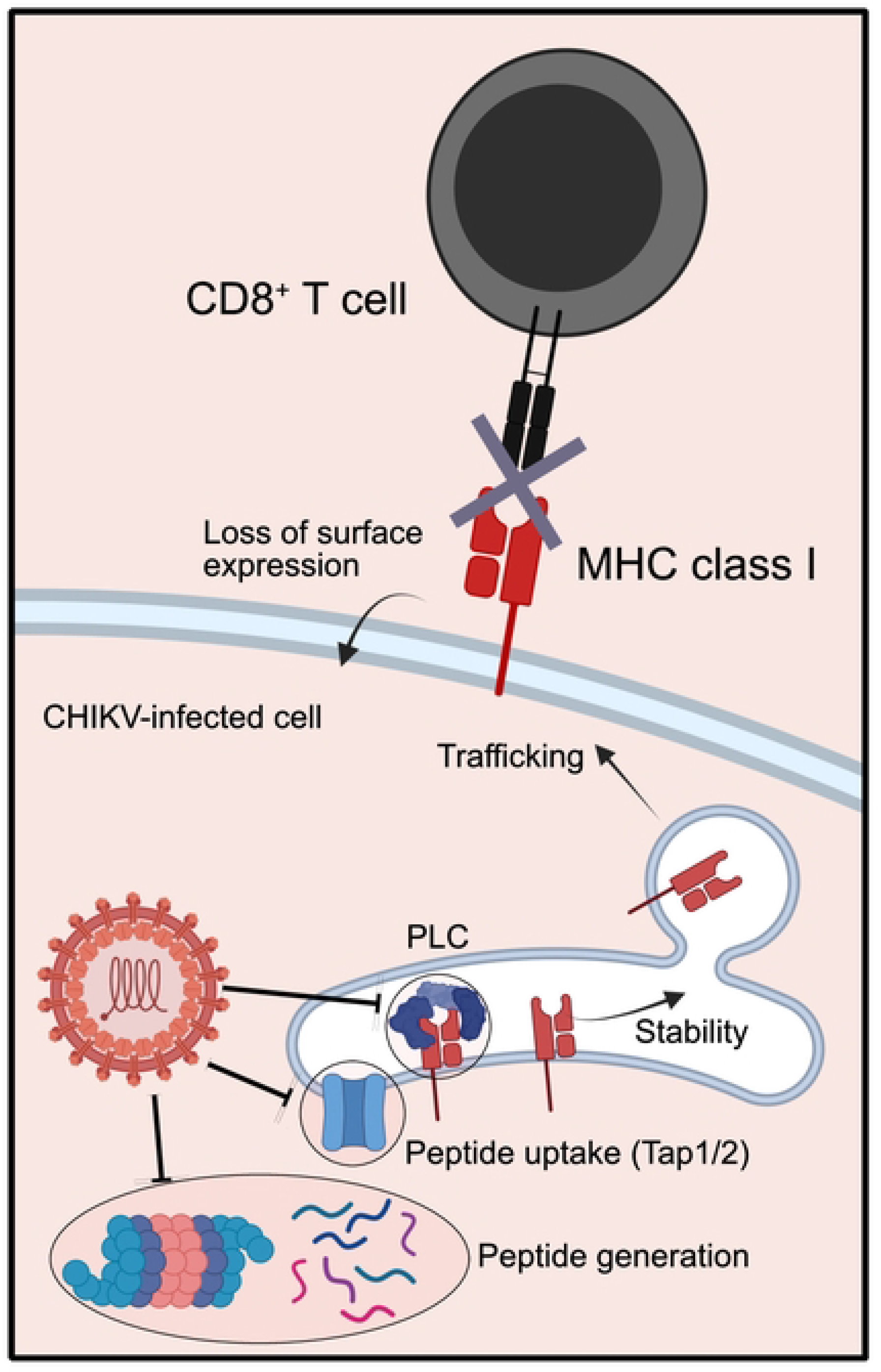
Proposed model for CHIKV disruption of MHC-I antigen presentation. CHIKV infection disables MHC-I antigen presentation at a step prior to β_2_m-peptide stabilization of the heavy chain within the ER. Figure made with Biorender.com.

## MATERIALS AND METHODS

### Ethics statement

This study was conducted in accordance with the recommendations in the Guide for the Care and Use of Laboratory Animals and the American Veterinary Medical Association (AVMA) Guidelines for the Euthanasia of Animals. All animal experiments conducted at the University of Colorado Anschutz Medical Campus were performed with the approval of the Institutional Animal Care and Use Committee (IACUC) of the University of Colorado School of Medicine (Assurance Number: A3269-01) under protocols 00026 and 00215. Experimental animals were humanely euthanized at defined endpoints by exposure to isoflurane vapors followed by bilateral thoracotomy.

### Mice and mouse experiments

An animal biosafety level 3 laboratory (ABSL3) with biosafety cabinet was used for all mouse experiments. C57BL/6 mice (stock #000664) were obtained from The Jackson Laboratory. CD45.1^+^ OT-I (C57BL/6-Tg(TcraTcrb)1100Mjb/J) mice were bred and housed at the University of Colorado School of Medicine under specific pathogen-free conditions. CD45.1^+^ OT-1 mice were maintained as hemizygous animals and selected based on flow cytometry phenotyping of transgene positive CD8^+^ cells expressing both Vα2 and Vβ5.1/2 TCRs. All mice were used in experiments at 4 weeks of age. Mice were anesthetized with isoflurane vapors and subcutaneously inoculated with 10^3^ PFU of CHIKV in the left rear footpad in 10 μL of PBS/1% FBS using a Hamilton syringe and 22G needle. Mice were euthanized by exposure to isoflurane vapors and bilateral thoracotomy, and then intracardially perfused with phosphate buffered saline (PBS) at the indicated time points.

### Cells

BHK-21 cells (American Type Culture Collection, ATCC CCL-10) were cultured at 37°C in Minimum Essential Medium Alpha (MEMα) supplemented with 10% fetal bovine serum (FBS), 10% tryptose phosphate broth (TPB) and 100 U/mL of penicillin-streptomycin (P/S). Mouse embryonic fibroblasts (derived from WT and *Ifnar1*^-/-^ C57BL/6 mice) were a gift from Michael Diamond (Washington University at St. Louis) and cultured in Dulbecco’s Modified Eagle medium (DMEM; HyClone 11965-084) supplemented with 10% FBS and 100 U/ml P/S (D:10). All cell lines were passaged when the monolayer was <90% confluent and fewer than 10 times. To generate primary *ex vivo* ankle (EVA) cells, 4-week-old C57BL/6 mice were euthanized, and ankle tissues dissected. A single-cell suspension was generated by horizontal shaking of ankle tissue dissections at 37°C for 2 h with 5 mm glass beads in digestion media containing D:10, 2.5 mg/mL collagenase type I (Worthington Biochemical), and 1.7 mg/mL DNase I (Roche). The cell suspension was filtered through a 100 μm sterile filter, washed with 1x PBS and centrifuged at 350 x g for 5 min at room temperature. The supernatant was decanted, and the cell pellet was resuspended in D:10. Approximately 1.5 x 10^6^ ankle tissue cells were added to a single well of a 6-well plate in 4 mL of D:10 medium and cultured at 37°C for 24 h. For virus infection studies, medium was aspirated, and cells were inoculated with 1 x 10^6^ PFU of indicated virus in 200 μl of *in vitro* diluent (PBS with 1% FBS, 1 mM Ca^2+^, and 1 mM Mg^2+^). After one hour with gentle rocking, the inoculum was removed, cells were washed with 1x PBS and then incubated for 16 h at 37°C in 4 mL D:10.

### Viruses

CHIKV cDNA clones of AF15561 were generated as described previously (67, 84). The recombinant CHIKV-VENKL cDNA clone was constructed by insertion of a tandem sequence encoding the fluorescent GFP derivative, VENUS, linked to LELQE-SIINFEKL-TEW followed by the 2A protease sequence of foot-and-mouth disease virus (FMDV2a) and inserted in-frame between capsid and E3 of CHIKV (45). CHIKV-SIINFEKLβ_2_m was generated similarly by inserting SIINFEKL with a GSSGSSGS linker between the β_2_m signal sequence and transcriptional start site of human β_2_m. CHIKV^QMS^-VENKL was generated by site-directed mutagenesis of the sequence encoding amino acids _674_ATL_677_ of the nsP2 using primers 5’ CACTAGGTCATACCTACCACTCATCTGTGGTAGACCCAgCTCTAGG-3’ and 5’-CCTAGAGCTGGGTCTACCACAGATGAGTGGTAGGTATGACCTAGTG-3’. The PCR amplicon was inserted into the WT vector. The complete viral genome sequence in all plasmids was confirmed by Sanger sequencing. Stocks of infectious CHIKV were generated from cDNA clones and titered by plaque assay on BHK-21 cells (67) or focus formation assay using Vero cells as previously described (85).

### *In vitro* virus replication assay

Triplicate wells of MEFs were inoculated with virus at a multiplicity of infection (MOI) of 0.01 FFU/cell. Viruses were absorbed onto cells for 1 h at 37°C with gentle rocking. Cells were then washed two times with 1 mL of 1x PBS, then 1 mL of D:10 was then added to each well, and cells were incubated at 37°C. At the indicated time points, 100 μL samples of culture supernatants were collected, and an equal volume of D:10 was added to maintain a constant volume within each well. Samples were stored at −80°C before analysis by focus formation assay on BHK-21 cells.

### Focus formation assays

As previously described (85), Vero cells were seeded in 96-well plates and inoculated with 10-fold serial dilutions of samples for 2 h at 37°C. Cells were overlayed with 0.5% methylcellulose in MEM-alpha plus 10% FBS and incubated for 18 h at 37°C. Following fixation with 1% paraformaldehyde (PFA), CHIKV-infected cells were detected using CHK-11 monoclonal antibody (86) at 500 ng/mL, and foci were visualized with TrueBlue Peroxidase substrate (SeraCare 5510–0030) and counted using a CTL Biospot analyzer and Biospot software (Cellular Technology).

### Flow cytometry

Single cell suspensions were generated either by trypsinizing cell monolayers from *in vitro* cultures or releasing cells from ankle tissue by mechanical and enzymatic digestion as previously described (41). Single cell suspensions were blocked with anti-FcγRIII/II (2.4G2; BD Pharmingen) for 10 min at RT, stained with LIVE/DEAD Fixable Violet Dead Cell Stain (ThermoFisher) according to product instructions, then stained with indicated antibodies for 45 min on ice. Cells were washed 2x with FACS buffer (1% FBS, 2 uM EDTA, 20 mM HEPES in 1x PBS) and fixed by addition of 1% paraformaldehyde for 10 min at RT. For intracellular stains, fixed cells were incubated with addition of indicated antibodies in 0.1% saponin + FACS buffer for 0.5-2 h at RT, washed 3x with 0.1% saponin + FACS buffer and resuspended in FACS buffer. Samples were acquired on a BD LSR Fortessa cytometer using FACSDiva software, or Cytek Aurora using Aurora software. Downstream analysis was performed using FlowJo software (Tree Star). Antibodies used: CD45 (30-F11), H2-Kb (AF6-88.5), H2-Kb-SIINFEKL (25D1.16), CD29 (TS2/16), CD44 (IM7) H2-Db (KH95), TCR (Vb8.2/8.3), TCR (Va2), CD8 (53-6.7), IL2Ra (7D4/CD25), IRF4 (3E4) and CD69 (H1.2F3).

### Western blotting

EVA Cells were trypsinized, washed twice with PBS then resuspended in cold NP-40 lysis buffer (1% NP-40, 50mM Tris-Cl, 150mM NaCl) with protease inhibitors (Cell Signaling, #5871) on ice for 30 min. Debris was removed by centrifugation at 17,000 g and assessed for total protein content by Bradford assay (ThermoFisher, A55866). 10 ug of lysate was denatured by addition of 5% 2-mercaptoethanol and 0.1% SDS-containing sample buffer and heating at 95°C for 5 min. Samples were ran via 8% SDS-PAGE, transferred onto polyvinylidene difluoride (PVDF) membranes by wet transfer and blocked in 10% milk in TBS containing 0.1% Tween (TBST). Anti-rabbit nsP2 (HLA1488, GeneTex) was used 1:500 overnight at 4°C in 5% milk TBST. Anti-mouse actin (mAbGEa, ThermoFisher) was used 1:5000 1 h RT in 5% milk TBST. After washing in TBST, secondary antibodies, goat anti-mouse IR700 (Azurebio AC2135) and anti-rabbit IR800 (Azurebio AC2128), were used 1:20,000 1 h at RT in 5% milk TBST. After washing in TBST, blots were imaged using Azure spec 500 NIR fluorescent imager.

### CD8^+^ T cell activation assays

Splenocytes from CD45.1^+^ OT-I mice were acquired as previously described (41). CD8^+^ T cells were enriched by magnetic bead separation with negative selection following the StemCell protocol (StemCell, 19853) and only utilized with >90% purity. After enrichment, CD45.1^+^ OT-I CD8^+^ T cells were added to primary murine ankle cell cultures that had been previously infected for 24h. Non-adherent cells were collected, stained with designated antibodies after 6 h of co-culture and analyzed by flow cytometry.

### Quantification and statistical analysis

Antigen presentation efficiency was calculated utilizing the Flowjo “derive parameter” function taking the mean fluorescence intensity of H2-K^b^-SIINFEKL channel and dividing it by VENUS or E2 mean fluorescence intensity (46). For statistical analysis, one-way analysis of variance (ANOVA) with multiple comparisons, and student’s t-test were performed on data using GraphPad Prism 10.02. P values of <0.05 were considered significant.

## ACKNOWLEDGMENTS

We thank the University of Colorado Anschutz Medical Campus Cancer Center Flow Cytometry Shared Resource Core Facility (RRID:SCR_022035). We thank Michael S. Diamond (Washington University) for critical evaluation of the manuscript. Finally, deep thanks to OP for technical support and assistance.

## SUPPORTING INFORMATION

**S1 Figure. Gating strategy for CHIKV-VENKL vs MOCK infection of EVA cells and correlation between MHC-I expression and productive infection.** (**A-C**) EVA cells were mock-inoculated or inoculated with 10^6^ PFU CHIKV-VENKL and assessed for VENUS and MHC-I cell surface expression at 24 hpi by flow cytometry. (**A**) Flow cytometry plots demonstrating gating strategy for VENUS^+^ or VENUS^-^ populations and CD29 expression. (**B-C**) Quantification and representative histograms of MHC-I cell surface expression stratified by VENUS expression [nil expression (grey), low (pink), medium (red), and high (dark red]. Data are representative from 2 independent experiments. P values were determined by one-way ANOVA with Tukey’s multiple comparison test. **, *P*<0.01; ***, *P*<0.001; ****, *P*<0.0001.

**S2 Figure. Gating strategy for CHIKV-VENKL- vs CHIKV^QMS^-VENKL-infected EVA cells.** Representative flow cytometry plots demonstrating gating strategy for live, CD45^-^, CD29^+^, VENUS^+^ fibroblasts from EVA cell cultures. Cell surface expression of CD29, H2-K^b^, H2-K^b^SIINFEKL and H2-D^b^ were assessed within live, singlets, CD45^-^, VENUS^+^ or VENUS^-^ gates.

**S3 Figure. OT-I CD8^+^ T cell enrichment efficiency, gating scheme, and baseline activation.** CD8^+^ T cells from naïve OT-I x CD45.1 mice were negatively selected for CD8^+^ cells by magnetic separation enrichment. (**A**) Frequency of CD8^+^ T cells before and after enrichment. (**B**) Gating scheme to differentiate bystander (K^b^OVA^257-264^ tetramer^-^) from OT-I (K^b^OVA^257-264^ tetramer^+^) CD8^+^ T cells. (**C**) Baseline activation of bystander (light grey; upper plot)) and OT-I (black; lower plot) CD8^+^ T cells without coculture with EVA cells.

**S4 Figure. Gating strategy for CHIKV-VENKL vs CHIKV^QMS^-VENKL infected ex vivo joint-associated fibroblasts**. Representative flow cytometry plots illustrating the gating strategy for live, CD45^-^, CD29^+^, VENUS^+^ fibroblasts from ex vivo joint-associated tissues. Cell surface expression of CD29, CD44, H2-K^b^, and H2-K^b^SIINFEKL were assessed within live, singlet, CD45^-^, VENUS^+^ or VENUS^-^ gates.

**S5 Figure. E2 and VENUS are equally detectable in CHIKV-infected EVA cells at 24 hpi.** EVA cells were inoculated with 10^6^ PFU of CHIKV-VENKL for 24hpi, harvested, fixed, permeabilized, and stained with anti-E2 CHIKV (CHK-11) at decreasing (1:2) serial dilutions. Cells were analyzed for E2 and VENUS expression by flow cytometry. (**A**) Representative flow cytometry plots. (**B**) Quantification of cell frequency for E2 and VENUS positive cells (red= infected, black = mock).

